# Three-dimensional anatomy of the early Eocene *Whitephippus* (Teleostei: Lampriformes) documents parallel conquests of the pelagic environment by multiple teleost lineages

**DOI:** 10.1101/2023.06.13.544748

**Authors:** Donald Davesne, James V. Andrews, Hermione T. Beckett, Sam Giles, Matt Friedman

**Author notes:** Corresponding authors: Donald Davesne, James V. Andrews. DD and JVA contributed equally to this work.

## Abstract

The early Eocene fossil assemblage of the London Clay (Southeastern England) is a key window to the early Paleogene diversification of teleost fishes in the open ocean. Despite their three-dimensional preservation that offers unique insight into skeletal anatomy, the London Clay fossils are still poorly described for the most part. †*Whitephippus tamensis* is a fossil teleost from this assemblage, known by several well-preserved specimens. Based on a complete description of the known material, including previously hidden structures (braincase, hyoid and branchial arches) revealed through 3D microtomography, we reinterpret †*Whitephippus* as an early member of the teleost group Lampriformes. More specifically, the anatomy of †*Whitephippus* indicates that it is likely a member of the so-called ‘pelagic clade’ including modern opahs and oarfishes. This redescription of †*Whitephippus* provides the earliest definitive evidence of lampriforms conquering the pelagic environment, alongside numerous other teleost lineages.

## INTRODUCTION

Actinopterygians (ray-finned fishes) are the primary vertebrate component of open-ocean ecosystems. The Cretaceous–Paleogene extinction event shaped this modern diversity by selectively affecting large predatory pelagic actinopterygian taxa, such as pachycormiforms, ichthyodectiforms, pachyrhizodontids and large aulopiforms (Cavin, 2001; Friedman, 2009; Friedman & Sallan, 2012). Their ecological equivalents in the modern ocean are mostly spiny-rayed teleosts (acanthomorphs) such as scombrids (e.g., tunas), xiphioids (billfishes) and carangoids (e.g., jacks and dolphinfishes). Although they all represent distinct evolutionary lineages, these groups of fast-swimming predators share gross similarities like streamlined bodies and falcate caudal fins.

Evidence from the fossil record (Friedman, 2010; Friedman et al., 2023; Guinot & Cavin, 2016) and node-dated molecular studies (Alfaro et al., 2009, 2018; Collar et al., 2022; Friedman et al., 2019; Ghezelayagh et al., 2022; Harrington et al., 2016; Miya et al., 2013) suggest that acanthomorph lineage diversity and body shape disparity exploded in the early Paleogene. Pelagic taxa in particular have been proposed to represent a case of parallel adaptive radiations, triggered by the conquest of ecological niches left vacant by the Cretaceous–Paleogene extinction event (Friedman, 2010; Miya et al., 2013). The most prominent groups of pelagic predatory acanthomorphs have their earliest fossil appearances clustered in a relatively short time interval in the late Paleocene–early Eocene (Fierstine, 2006; Monsch & Bannikov, 2011; Santini et al., 2013; Santini & Carnevale, 2015). However, a relatively low number of fossil sites document teleost diversity in the open ocean during this key time interval (Argyriou & Davesne, 2021; El-Sayed et al., 2021; Friedman et al., 2016).

The London Clay of Southeastern England is one of these fossil localities. This Ypresian (early Eocene, c. 52–49 Ma) assemblage preserves a very diverse marine teleost fauna with very few equivalents in the early Paleogene in terms of taxonomic richness (Casier, 1966; Friedman et al., 2016). Its composition closely matches that of modern open ocean faunas, with taxa like scombrids (Beckett & Friedman, 2016; Monsch, 2005), xiphioids (Monsch, 2005), luvarids (Bannikov & Tyler, 1995), megalopids (Forey, 1973), trichiuroids (Beckett et al., 2018) and carangids (Casier, 1966; Friedman et al., 2016; Monsch, 2005). Remarkably, teleosts from the London Clay are preserved in three dimensions, with minimal deformation and fine internal and external anatomical details preserved in life position. These unique taphonomic conditions offer the possibility to study phylogenetically informative anatomical structures that would usually be concealed in flattened fossils, such as the braincase and branchial arches (e.g. Beckett and Friedman, 2016; Close et al., 2016; Friedman et al., 2016; Beckett et al., 2018). The diversity of the London Clay fauna has been noted as early as the 19^th^ century (e.g., Agassiz, 1845), described by Woodward (1901) and later revised in detail by Casier (1966). However, some of the taxa referred to in these publications are probably misclassified (Patterson, 1993), and the phylogenetic positions of most are poorly constrained (Friedman et al., 2016).

†*Whitephippus* is one such case. This genus, known by several specimens (Figs. 1–3, 7, 9), has been attributed to the extant family Ephippidae (spadefishes) by Woodward (1901) and Casier (1966). However, some authors (Bonde, 1995; Carnevale, 2004; Friedman et al., 2016) have proposed that †*Whitephippus* should be classified with Lampriformes instead. Lampriformes are a clade of marine pelagic acanthomorphs that include such distinctive taxa as the endothermic opah (Lampridae) and the giant elongated oarfish (Regalecidae). Lampriform fossil taxa are relatively numerous, documenting the extreme morphological changes underwent in the early evolution of the group (Bannikov, 1999, 2014; Davesne, 2017; Davesne et al., 2014). Significantly, †*Whitephippus* is the only fossil lampriform known by complete and three-dimensionally preserved skulls (Figs. 1–3). Using computed microtomography (μCT), we reevaluate the anatomy and phylogenetic position of †*Whitephippus*, giving a unique perspective of the evolution of lampriforms in the context of the origin of modern pelagic actinopterygian faunas.

**Figure 1.**
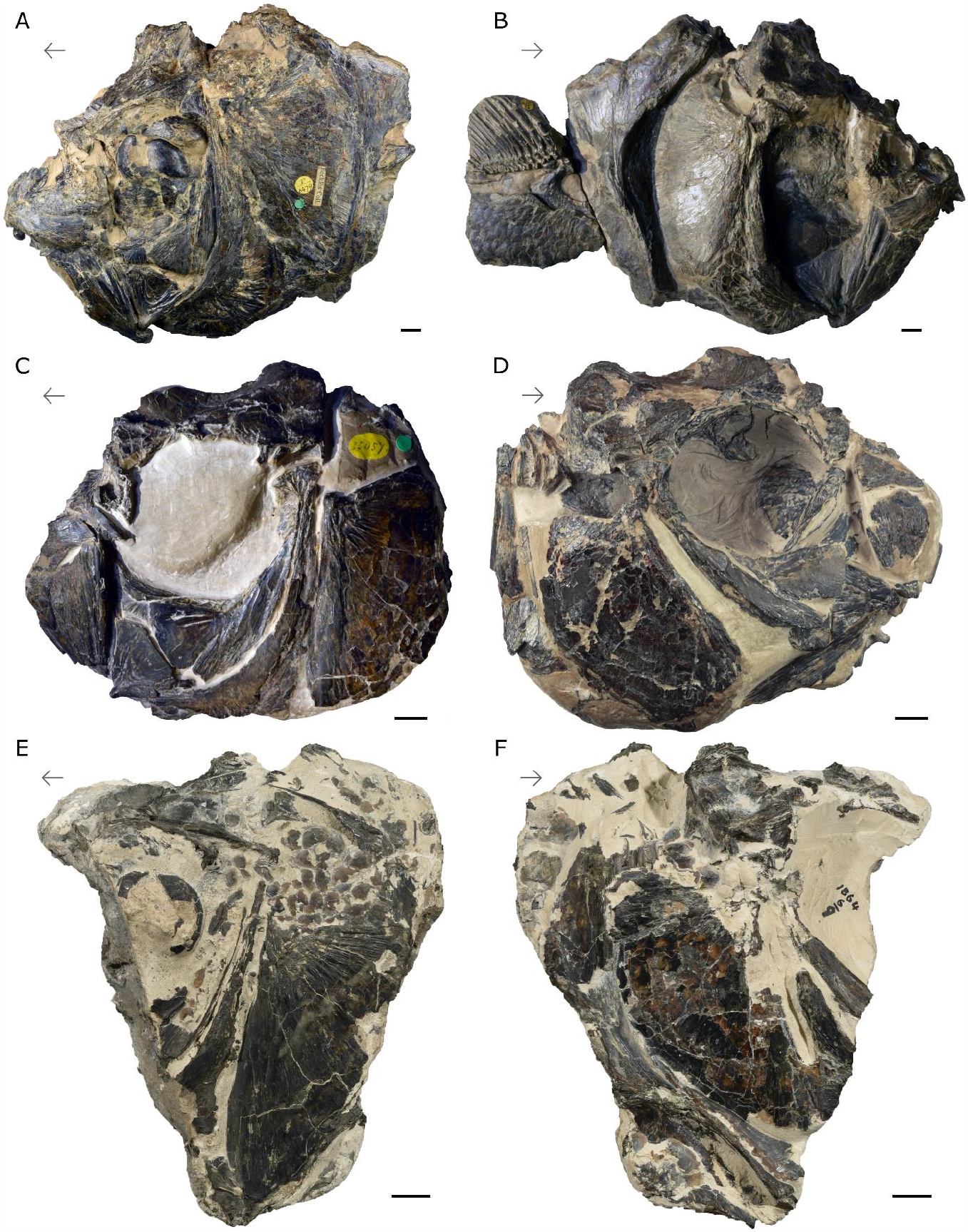
†*Whitephippus tamensis*. London Clay Formation, early Eocene (Ypresian), Southeast England, UK. Photographs of holotype NHMUK PV P 6479 in (**A**) left and (**B**) right lateral views; †*Whitephippus* cf. *tamensis* NHMUK PV P 35057 in (**C**) left and (**D**) right lateral views; NMS 1864.6.9 in (**E**) left and (**F**) right lateral views. Arrows indicate anatomical anterior. Scale bars represent 1 cm.

## Material and Methods

### Specimen imaging

Computed tomography. NHMUK fossil specimens were imaged using the Nikon XT H 225 ST industrial CT scanner at the Natural History Museum, London. Reconstructed datasets were segmented and visualized in Mimics v 19.0 (Materialise, Belgium). A fossil specimen from NMS and a recent specimen from UMMZ were imaged using the same model of instrument in the CTEES facility at the University of Michigan. Osteological figures were generated in Blender v 2.91 (blender.org) from surface (.ply) files. The tomograms surface files of segmented elements are available on MorphoSource (www.morphosource.org/projects/000561545).

Institutional Abbreviations— Institutional abbreviations follow those of Sabaj (2020). **AMNH**, American Museum of Natural History, New York, NY, USA; **MNHN**, Muséum national d’Histoire naturelle, Paris, France; **NHMUK**, Natural History Museum, London, UK; **NMS**, National Museums of Scotland, Edinburgh, UK; **UMMZ**, University of Michigan Museum of Zoology, Ann Arbor, MI, USA; **ZMUC**, Statens Naturhistoriske Museum, Københavns Universitet, Copenhagen, Denmark.

Dagger symbol— The obelus (†) indicates extinct taxa, following Patterson & Rosen (1977).

Comparative material, fossil— †*Whitephippus tamensis*, NHMUK PV P 6479, holotype, almost complete cranium with parts of the pectoral girdle and fin, μCT-scanned (Figs. 1A-B, 9); †*Whitephippus tamensis*, NHMUK PV 41384, paratype, isolated posterior portion of the neurocranium (Supp. Fig. 1); †*Whitephippus* cf. *tamensis*, NHMUK PV OR 35057, almost complete cranium, μCT-scanned (Figs. 1C-D, 2A-B, 3A-F, 5A-C, 6A-B; Supp. Figs. 2A-B, 3); †*Whitephippus* cf. *tamensis*, NMS 1864.6.9, posterior portion of the cranium, vertebrae and pectoral girdle, μCT-scanned (Figs. 1E-F, 2C-D, 3G-L, 7; Supp. Fig. 2C-D).

**Figure 2.**
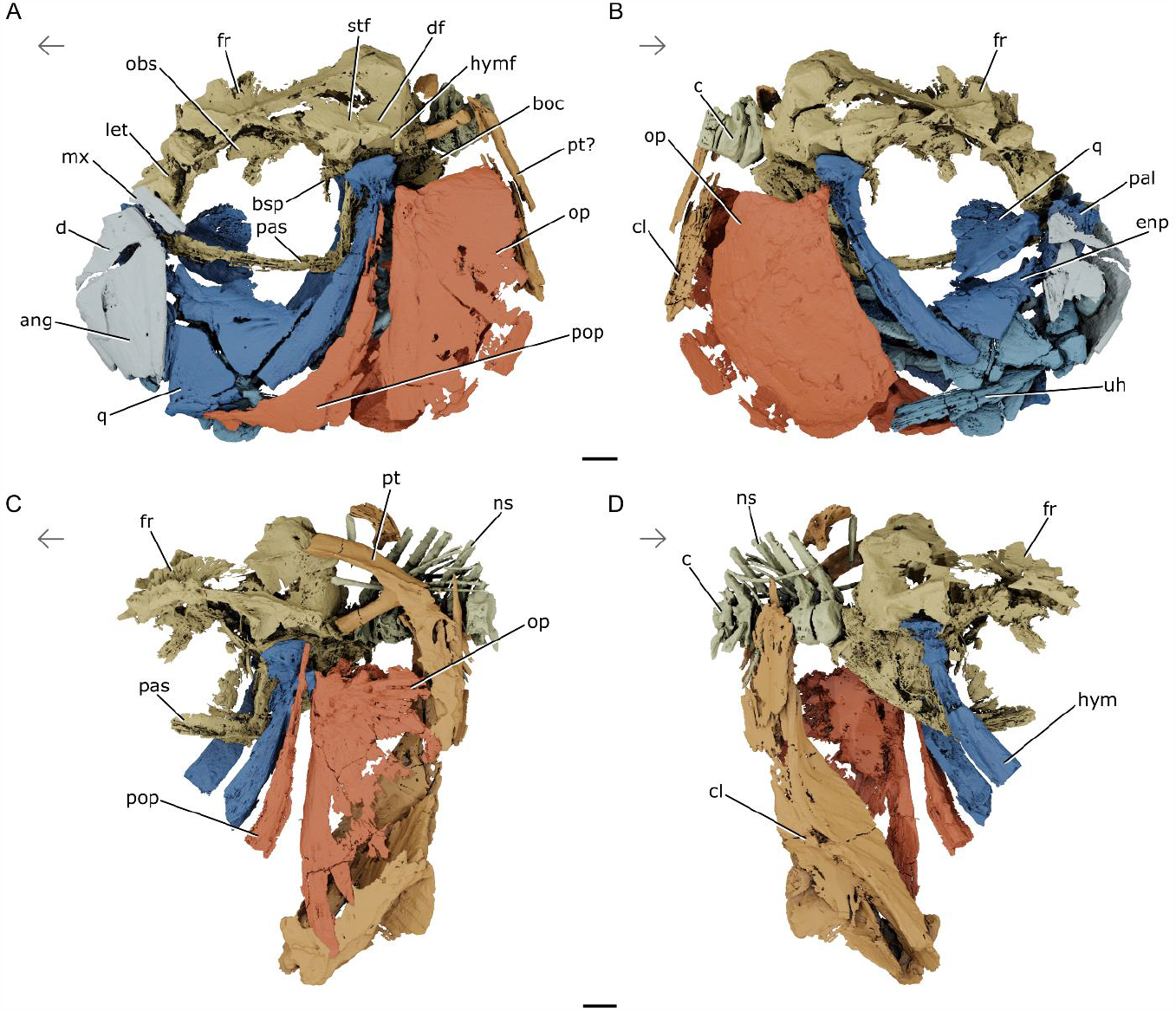
Skull, vertebrae, and pectoral girdle of †*Whitephippus* cf. *tamensis*. London Clay Formation, early Eocene (Ypresian), Southeast England, UK. Rendered μCT models showing NHMUK PV P 35057 in (**A**) left and (**B**) right side views; NMS 1864.6.9 in (**C**) left and (**D**) right side views. Skeletal regions highlighted as follows: neurocranium (yellow), suspensorium (dark blue), jaws (platinum), opercles (sienna), ventral hyoid (blue), gill skeleton (light blue), pectoral girdle (orange), vertebral column (light yellow). Abbreviations: **ang**, anguloarticular; **boc**, basioccipital; **bsp**, basisphenoid; **c**, centra; **cl**, cleithrum; **d**, dentary; **df**, dilatator fossa; **enp**, endopterygoid; **fr**, frontal; **hym**, hyomandibula; **hymf**, hyomandibular facet; **let**, lateral ethmoid; **mx**, maxilla; **ns**, neural spine; **obs**, orbitosphenoid; **op**, opercle; **pal**, palatine; **pas**, parasphenoid; **pop**, preopercle; **pt**, posttemporal; **q**, quadrate; **stf**, supratemporal fossa; **uh**, urohyal. Arrows indicate anatomical anterior. Scale bars represent 1 cm.

**Figure 3.**
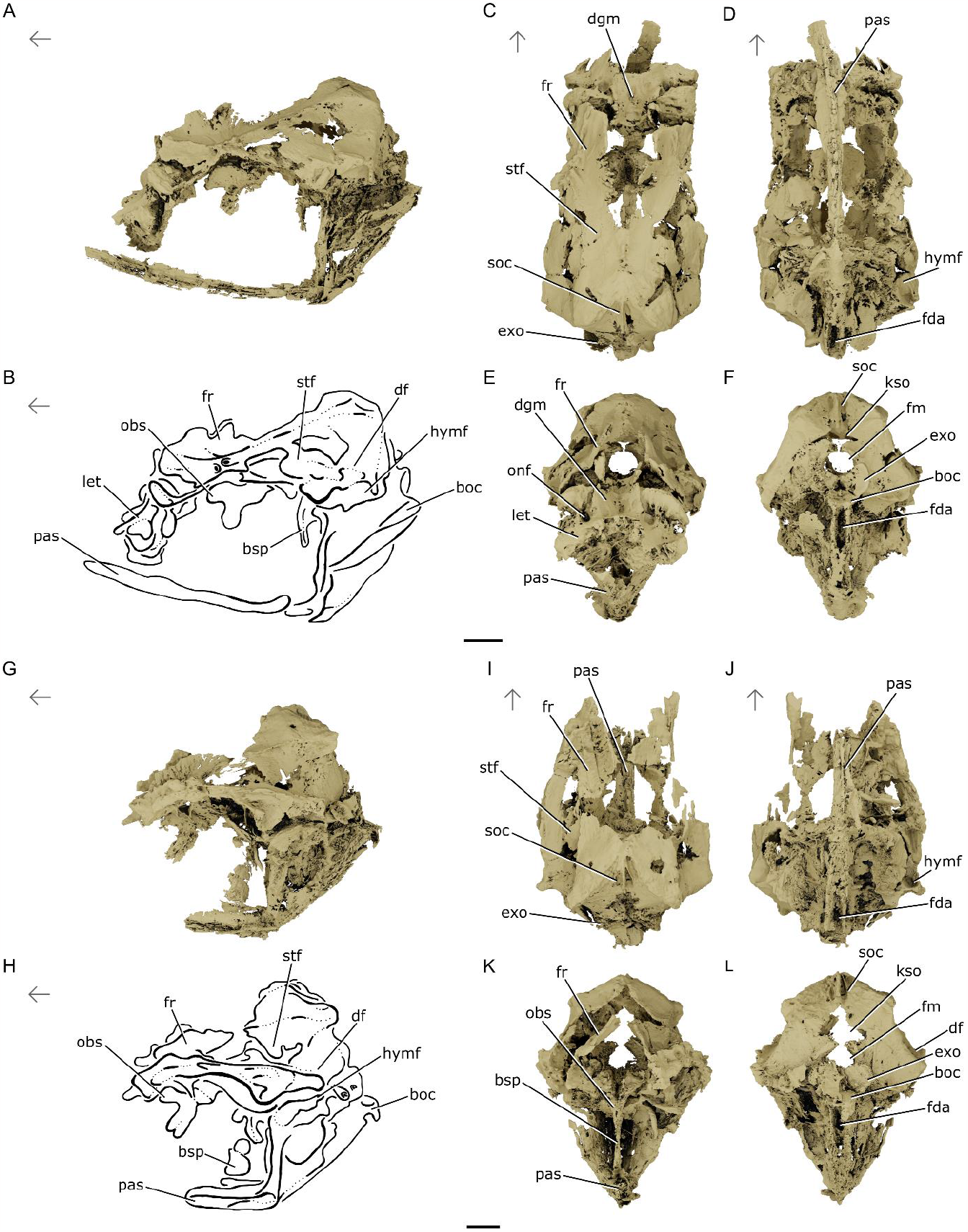
Neurocranium of †*Whitephippus* cf. *tamensis*. London Clay Formation, early Eocene (Ypresian), Southeast England, UK. Rendered μCT models and line drawings showing NHMUK PV P 35057 in (**A & B**) lateral, (**C**) dorsal, (**D**) ventral, (**E**) anterior, and (**F**) posterior views; NMS 1864.6.9 in (**G & H**) lateral, (**I**) dorsal, (**J**) ventral, (**K**) anterior, and (**L**) posterior views Abbreviations: **boc**, basioccipital; **bsp**, basisphenoid; **df**, dilatator fossa; **dgm**, dorsal groove of mesethmoid; **exo**, exoccipital; **fda**, fossa for dorsal aorta; **fm**, foramen magnum; **fr**, frontal; **hymf**, hyomandibular facet; **kso**, “keyhole-shaped opening”; **let**, lateral ethmoid; **obs**, orbitosphenoid; **onf**, olfactory nerve foramen; **pas**, parasphenoid; **soc**, supraoccipital; **stf**, supratemporal fossa. Arrows indicate anatomical anterior. Scale bars represent 1 cm.

Comparative material, extant— *Lampris* cf. *guttatus*, AMNH 79669 SD (Figs. 4F-I, 5G-H, 6D, 8B, E-F), AMNH 21720 SD (Supp. Figs. 4-6), MNHN.ZA.1883-1795, ZMUC 74 (dry osteological preparations; specific attribution follows that accompanying these materials, and we acknowledge that these might belong to other species of *Lampris*; Underkoffler et al., 2018); *Metavelifer multiradiatus*, AMNH 214663 SD, 219280 SD, 91808 SD, 91800 SD and 91798 SD (dry osteological preparations); *Velifer hypselopterus*, MNHN.IC.1982.0025, UMMZ 220456 (preserved in alcohol and μCT-scanned; Figs. 4A-E, 5D-F, 6C, 8A, C-D).

**Figure 4.**
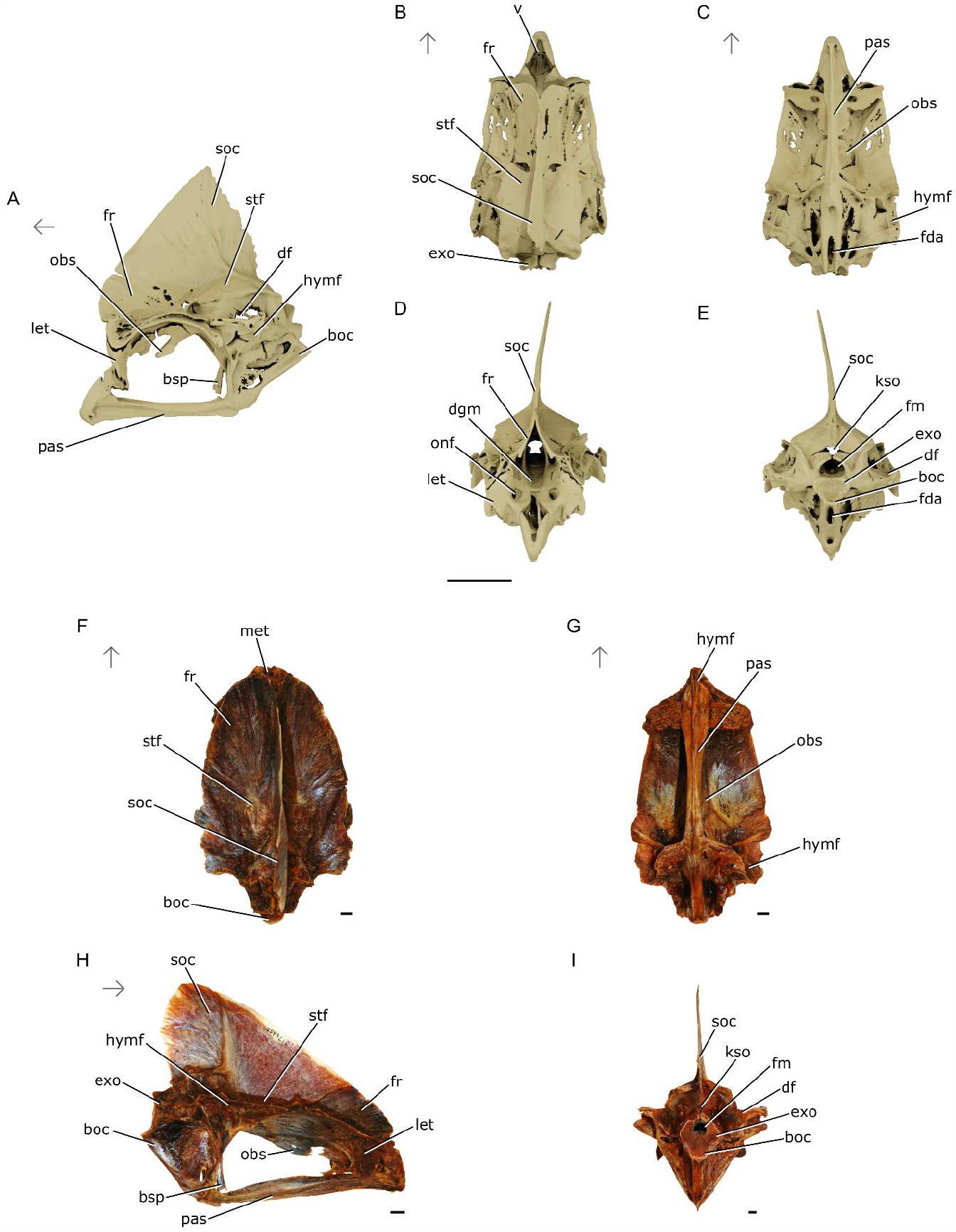
Neurocrania of relevant extant lampriforms. Rendered μCT models of *Velifer hypselopterus* UMMZ 220456 showing the (**A**) lateral, (**B**) dorsal, (**C**) ventral, (**D**) anterior, and (**E**) posterior views. Photographs of *Lampris* cf. *guttatus* AMNH 79669SD in (**F**) dorsal, (**G**) ventral, (**H**) lateral, and (**I**) posterior views. Abbreviations: **boc**, basioccipital; **bsp**, basisphenoid; **df**, dilatator fossa; **dgm**, dorsal groove of mesethmoid; **exo**, exoccipital; **fda**, fossa for dorsal aorta; **fm**, foramen magnum; **fr**, frontal; **hymf**, hyomandibular facet; **kso**, “keyhole-shaped opening”; **let**, lateral ethmoid; **met**, mesethmoid; **obs**, orbitosphenoid; **onf**, olfactory nerve foramen; **pas**, parasphenoid; **soc**, supraoccipital; **stf**, supratempora fossa; **v**, vomer. Arrows indicate anatomical anterior. Scale bars represent 1 cm.

**Figure 5.**
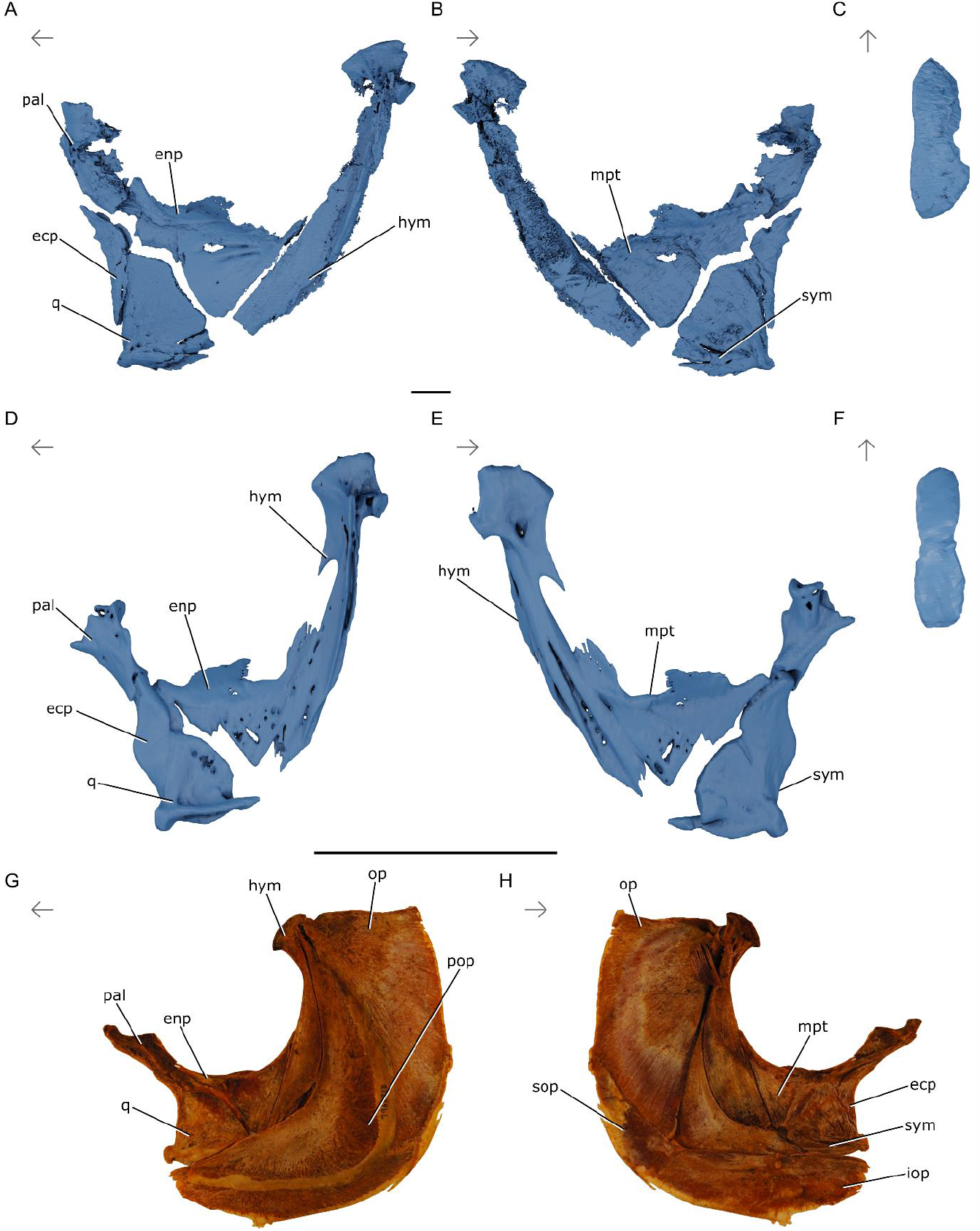
Left suspensoria of relevant lampriforms. †*Whitephippus* cf. *tamensis* NHMUK PV P 35057, London Clay Formation, early Eocene (Ypresian), Southeast England, UK; rendered μCT models showing (**A**) lateral and (**B**) mesial views, and (**C**) the hyomandibular head in proximal view; *Velifer hypselopterus* UMMZ 220456, rendered μCT models in (**D**) lateral and (**E**) mesial views, and (**F**) the hyomandibular head in proximal view; *Lampris* cf. *guttatus* AMNH 79669SD, photographs in (**G**) lateral and (**H**) mesial views. Abbreviations: **ecp**, ectopterygoid; **enp**, endopterygoid; **hym**, hyomandibula; **iop**, interopercle; **mpt**, metapterygoid; **op**, opercle; **pal**, palatine; **pop**, preopercle; **q**, quadrate; **sop**, subopercle; **sym**, symplectic. Arrows indicate anatomical anterior. Scale bars represent 1 cm. Hyomandibular heads not to scale.

**Figure 6.**
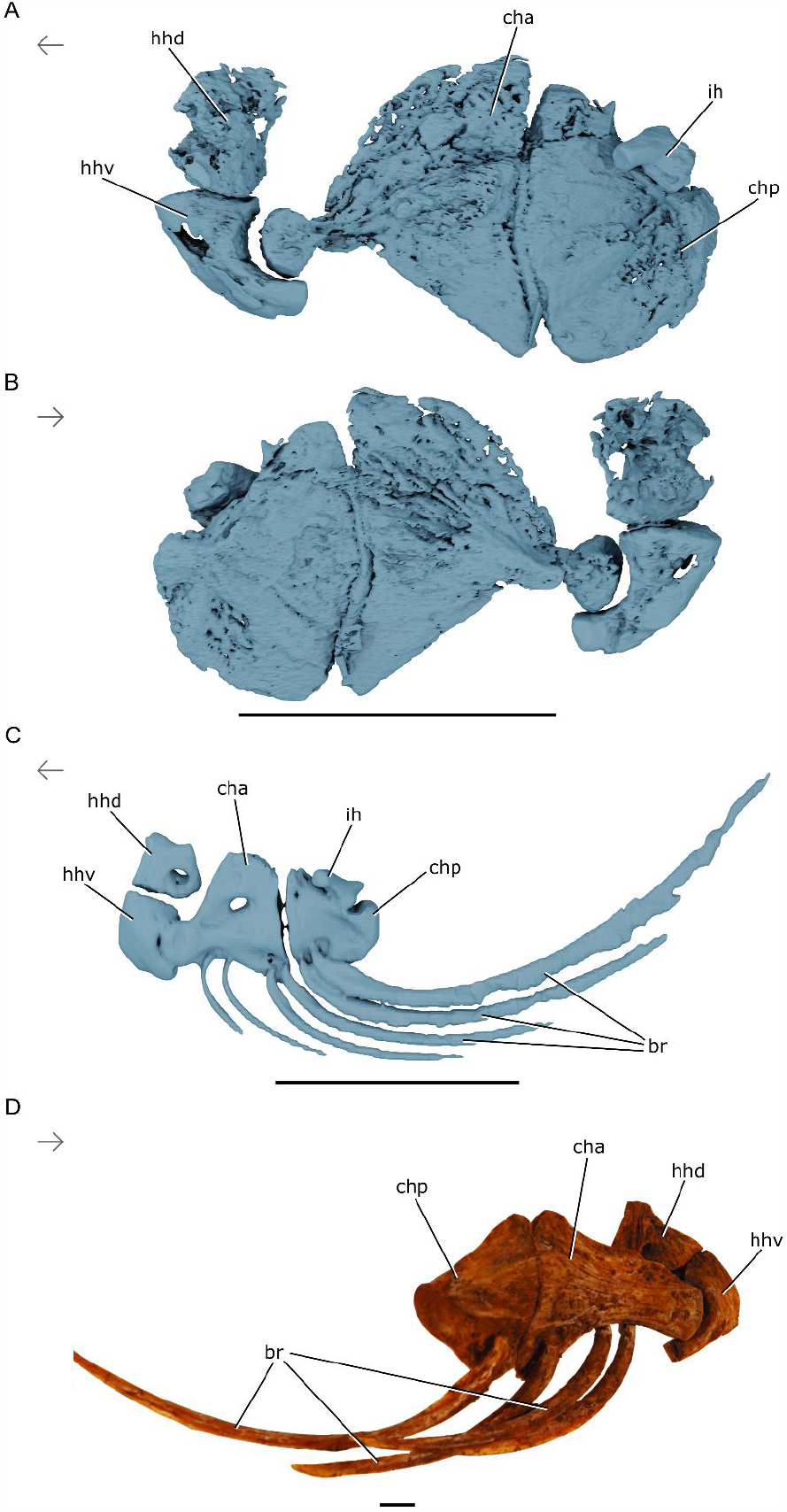
Ventral hyoids of relevant lampriforms. †*Whitephippus* cf. *tamensis* NHMUK PV P 35057, London Clay Formation, early Eocene (Ypresian), Southeast England, UK, rendered μCT models showing the left ventral hyoid in (**A**) lateral and (**B**) mesial views; *Velifer hypselopterus* UMMZ 220456, rendered μCT models of left ventral hyoid in (**C**) lateral view; *Lampris guttatus* AMNH 79669SD, photograph of right ventral hyoid in (**D**) lateral view. Abbreviations: **br**, branchiostegals; **cha**, anterior ceratohyal; **chp**, posterior ceratohyal; **hhd**, dorsal hypohyal; **hhv**, ventral hypohyal; **ih**, interhyal. Arrows indicate anatomical anterior. Scale bars represent 1 cm. Urohyal not imaged.

## SYSTEMATIC PALAEONTOLOGY

TELEOSTEI Müller, 1845

ACANTHOMORPHA Rosen, 1973

Order LAMPRIFORMES Goodrich, 1909

Genus †*WHITEPHIPPUS* Casier, 1966

†*Whitephippus tamensis* Casier, 1966

1901 †*Laparus alticeps* Woodward, p. 596

1966 †*Whitephippus tamensis* Casier, p. 237-243, figs. 52-54, pl. 31

*Holotype*. NHMUK PV P 6479.

*Paratype*. NHMUK PV 41384.

*Type locality*. Isle of Sheppey, Kent, England, United Kingdom.

*Stratigraphic age*. London Clay Formation, unit D (King, 1981, 1984). Ypresian, early Eocene, c. 52-49 Ma.

## Description

### Neurocranium

No specimen of †*Whitephippus* preserves a complete neurocranium. NHMUK PV 41384 represents an isolated neurocranium that is incomplete and largely free of surrounding matrix (Supp. Fig. 1). The neurocranium is incompletely exposed in NHMUK PV P6479, NHMUK PV 35057, and NMS 1864.6.9 (Fig. 1), but μCT reveals considerable detail of concealed anatomy in the latter two specimens (Figs. 2, 3). Our description of the neurocranium draws on all four specimens.

Anteriorly, the neurocranium includes a mesethmoid and paired lateral ethmoids that are tightly bound to one another (Fig. 3A-D). The junction between these bones is clear dorsally, where they are separated by a conspicuous vertical suture; they cannot be distinguished from one another more ventrally. The lateral ethmoid defines the anterior margin of the orbit, and its anterior and posterior surfaces are concave. The foramen for the olfactory nerve pierces the lateral ethmoid in its ventral half, near to the bone’s junction with the mesethmoid. Viewed anteriorly, the outer margins of the lateral ethmoids are strongly excavated at the level of this canal. The anterolateral corner of the lateral ethmoid articulates with the lacrimal (infraorbital 1).

The dorsal surface of the mesethmoid forms a trough (Fig. 3C). The mesethmoid extends slightly posterior to the lateral ethmoids when viewed laterally. The broad dorsal groove of the mesethmoid forms the floor of an open chamber, defined laterally by the broken edge of the frontals. Viewed dorsally, this chamber is ‘V’ shaped, with its apex pointing posteriorly. The incomplete frontals are flanged dorsally over their anterior third, such that their exterior surfaces face laterally. We interpret this arrangement as representing a broken frontal ‘vault’ that would have opened anteriorly but been enclosed laterally and dorsally. This vault accommodates the greatly elongated ascending processes of the premaxillae in extant lampriforms.

The divisions between the frontals and other bones are not prominent externally, and our μCT data are insufficient to resolve most of the sutures between ossifications of the neurocranium (Fig. 3). We therefore describe the gross morphology of the neurocranium, drawing attention to divisions between bones when these are visible. Posterior to the incomplete frontal vault, the roof of the skull is complete, and forms the broken base of a probable sagittal crest that extends along the posterior half of the frontal. Posterior to the frontal vault, the roof of the neurocranium is elevated relative to the ethmoid region, giving it a dome-like appearance in lateral view. The dorsolateral surface of the posterior half of the neurocranium is dominated by the supratemporal fossa (Fig. 3A, G). This shallow, triangular depression extends anteriorly to the level of mid-orbit. A fenestra lies in the centre of the supratemporal fossa. Sutures are visible extending from this fenestra. These define the edges of the bones contributing to the supratemporal fossa: the epiotic posterodorsally, the pterotic posterolaterally, the parietal anterodorsally, and the sphenotic anterolaterally. The dorsolateral surface of the epiotic bears a flattened facet for the posttemporal. A low ridge of bone defines the ventrolateral margin of the supratemporal fossa and separates it from the dilatator fossa. The narrow, oval-shaped hyomandibular facet lies ventral to the dilatator fossa and straddles the division between the pterotic and the sphenotic.

Viewed posteriorly, the braincase has a roughly triangular profile dorsal to the foramen magnum (Figs. 3F, L). The foramen magnum communicates dorsally with another opening in the back of the skull through a narrow gap between median extensions of the exoccipitals, forming an inverted keyhole-shaped opening. The supraoccipital extends posteroventrally to contribute to the dorsal fringe of this opening, but does not separate the exoccipitals themselves or contribute to the foramen magnum. The occipital condyle is located approximately at the level of the upper half of the orbit, placing it midway between the base of the neurocranium as defined by the parasphenoid and the dorsal peak of the supraoccipital as preserved. The occipital condyle is broadest dorsally, with large, kidney-shaped facets of the posterior margins of the exoccipitals that extend dorsolaterally to partially define the lateral margins of the foramen magnum (Fig. 3F, L; Supp. Fig. 1). These exoccipital condyles meet on the midline immediately ventral to the foramen magnum, but they are separated ventrally by the basioccipital condyle. This condyle has an angular dorsal margin and rounded ventral margin in posterior view and is roughly the size of a single exoccipital condyle.

Divisions between bones contributing to the ventrolateral walls of the otic region are not clear in the available material. In posterior view, these walls appear straight, joining ventrally in an acute angle that gives the braincase ventral to the occipital condyle the profile of an inverted triangle. In lateral view, the ventral margin of the otic and occipital regions is strongly sloped, defining an angle of roughly 45° with a line extending from the main axis of the parasphenoid in the orbital region. The posterior stalk of the parasphenoid approaches—but does not contact—the basioccipital condyle. A narrow fossa is present along the ventral margin of the basioccipital, and corresponds with where the dorsal aorta would have contacted the neurocranium in life (L. Grande & Bemis, 1998). It extends anteriorly towards the posterior stalk of the parasphenoid. The anterior and posterior portions of the parasphenoid join at a conspicuous angle at the level of the anterior margin of the myodome. The parasphenoid bears two sets of buttresses in this region, laterodorsal to the myodome and roughly perpendicular with the shaft of the parasphenoid. The more anterior of these is long and narrow, and extends along the anterior margin of the prootic, bracing the slender, rod-like lateral commissure. The posterior projection is shorter, but anteroposteriorly thickened, and is notched by a large foramen in the neurocranium for the internal carotid artery. Anterior to these processes, the parasphenoid widens in ventral view. There is a gap between the parasphenoid and the ethmoid region. We assume this would have been bridged by the vomer in life, but this bone is not preserved in available material.

The median basisphenoid is ‘T’ shaped in anterior view, and completely divides the optic foramen from the myodome (Fig. 3A-B, G-H). The ventral process of the basisphenoid is long, slender, and directed slightly posteriorly. The ventral portions of the basisphenoid in the NHMUK material are not clearly shown in μCT data, but NMS 1864.6.9 clearly shows a pedicel on the midline of the parasphenoid, reaching dorsally to nearly contact the basisphenoid (Fig. 3G). These bones were most likely separated by cartilage in life.

The optic foramen is large and situated on the posterodorsal roof of the interorbital region. It is delimited posteroventrally by the basisphenoid, laterally by the paired pterosphenoids, and anteriorly by the median orbitosphenoid (Fig. 3A, G). In ventral view, the orbitosphenoid is wider than long. Its posterior margin bears a deep notch where the bone delimits the optic foramen, and its anterior margin is more subtly excavated. The orbitosphenoid carries a pronounced keel along its ventral midline, which is directed at a subtle angle anteriorly and has a rounded ventral extremity.

### Infraorbitals, lacrimal and sclerotic ring

The lacrimal (infraorbital 1) is preserved in NHMUK PV P6479, but is highly fragmentary (Fig. 1B). It is a plate-like bone with a long anteroposterior axis. The bone is deepest posteriorly, and tapers to a rounded anterior apex. A process extends from the dorsal margin of the bone and bears a concave medial facet that articulates with the lateral ethmoid. The more posterior infraorbitals are not preserved.

Well-developed sclerotic ossicles are present in NHMUK PV P6479 and NMS 1864.6.9 (Fig. 1A, E).

### Jaws

The jaws are partially preserved in NHMUK PV P6479 and PV 35057 (Fig. 1A-D; Fig. 2A-B). The mandibles appear to have been oriented anterodorsally in life, similar to the configuration seen in extant *Velifer* and *Lampris*. The anguloarticular is high and triangular, being less tall near its articulation with the quadrate than at its contact with the dentary. A posteriorly oriented prong extends from the bone ventral to the glenoid fossa. In available material, a narrow gap separates the dentary from the anguloarticular. There is no autogenous retroarticular ossification visible in examined material. The dentary is incomplete in all available specimens. Anteriorly, it is rounded and appears edentulous, while the posterior portions show a straight dorsal margin with no visible teeth.

The upper jaws are partially preserved in NHMUK PV P6479 and PV 35057. They appear to be much smaller than the high and deep lower jaws. The maxilla is toothless and anteroposteriorly elongate (anterior fragment visible in Fig. 2A). Its posterior extremity is slightly rounded and it curves dorsally at its anterior end.

The ascending processes of the premaxillae are preserved in NHMUK PV P6479. They are closely associated together throughout their length. A bulbous pyritic mass lies between the ascending processes and protrudes anteriorly. This may correspond to a mineralized rostral cartilage, as Casier (1966) proposed. Apart from the ascending processes, no other parts of the premaxillae are visible due to a combination of breakage and concealment by overlying matrix and the maxillae.

### Suspensorium

A complete suspensorium is preserved in NHMUK PV 35057 (Figs. 2A, B; 5A-C). Only its ventral portion is visible externally, but concealed regions are clear in the μCT reconstructions.

The hyomandibula is very long and its ventral shaft is especially well developed. Its articular head is relatively narrow and directed slightly anteriorly at the level of its contact with the neurocranium. This zone of contact consists of a single condyle, slightly pinched dorsolaterally in its middle. Posteroventral to the articular head, a short condyle contacts the opercular bone, projecting slightly ventrally. A conspicuous lateral ridge extends from the articular head near or along the posterior edge of the ventral shaft of the hyomandibula. This ridge is approximately two-thirds the length of the entire ventral shaft and delimits an anteriorly-oriented fossa bounded by the anterior lamina of the ventral shaft of the hyomandibula. A narrow posterior lamina is restricted to the region immediately ventral to the opercular condyle of the hyomandibula and is completely covered by the preopercle. Anterior and ventral to the ridge, the surface of the hyomandibula is flat and faces laterally. Overall, the ventral shaft of the hyomandibula is strap-shaped, with a gentle convexity to its posterior margin. The ventral shaft is widest at the level of the dorsal articulation of the metapterygoid and terminates distally at a blunt tip, although this may be a taphonomic break.

The metapterygoid is a laminar bone that is roughly triangular in shape. The entire posterior edge of the bone contacts the anterior margin of the ventral shaft of the hyomandibula. The metapterygoid bears a posteriorly directed, spur-like process that extends from the posterodorsal corner of the bone, tracing the anterior margin of the hyomandibula. The principal surface of the metapterygoid faces laterally, but folds slightly medially along its dorsal edge.

The quadrate is subtriangular, with its tip forming a ventrally projecting condyle that articulates with the glenoid fossa of the anguloarticular. The posteroventral edge of the bone bears a thickening aligned with the quadrate condyle. The short symplectic rests in a notch that is aligned with this thickened ridge at the posterodorsal corner of the quadrate. The anterior margin of the quadrate bears a faint, laterally directed furrow at its contact with the ectopterygoid and anguloarticular. The dorsal margin of the bone comprises two edges, the more anterior of which forms a margin with the ectopterygoid, while the more posterior traces the profile of the anteroventral margin of the metapterygoid. In NHMUK PV 35057, the left quadrate is in life position, but the right one is displaced and upturned.

The endopterygoid is triradiate, with anterior, posterior and ventrolateral rami. The posterior and ventrolateral branches of the bone embrace the anterior angle of the metapterygoid. The flat surface of the posterior ramus shows a strong dorsomedial orientation. The anterior ramus bears a longitudinal ridge at its dorsal margin, presumably to strengthen the articulation with the ectopterygoid.

The anterior margin of the suspensorium is defined by the ectopterygoid and palatine. The ectopterygoid is slightly crescentic. Its posteroventral ramus is longer, narrower and more tapered than its anterior ramus; the two rami form a moderately concave ventral margin. The dorsal margin is more strongly pronounced and the mesial margin of the bone is slightly concave. The palatine is subrectangular. It contacts the ectopterygoid and endopterygoid posteriorly, and the braincase at the level of the lateral ethmoids anteriorly. There is no distinct palatine prong marking the articulation between the suspensorium and the upper jaw. However, the connection between the palatine and the braincase is marked by an anterodorsally directed articular head.

### Opercular series

Apart from the isolated braincase of NHMUK PV 41384, all specimens of †*Whitephippus* preserve portions of the opercle and preopercle (Figs. 1, 2). Portions of the interopercle are preserved in NHMUK PV P6479, where the bone is visible along the posteroventral margin of the preopercle. The opercle is deep, nearly reaching the ventral margin of the hyomandibular shaft. The anterior margin of the opercle is nearly vertical, while the dorsal margin is gently angled with its dorsal apex located at the posterior margin of the bone. The posterior margin of the bone is gently convex, and the opercle terminates ventrally in a blunt tip. The facet for the hyomandibula is located at the junction of the dorsal and anterior margins of the opercle and is produced as an anterodorsally extending projection in lateral view.

The apex of the dorsal limb of the preopercle is located immediately ventral to the articulation between the opercle and hyomandibula. The preopercle is crescent shaped, and the dorsal limb is substantially longer than the ventral one. At their extremities, these two limbs are oriented approximately perpendicular to one another.

### Branchial skeleton and ventral hyoid arch

The ventral hyoid arch and branchial skeleton are concealed in all specimens but are visible in tomograms of NHMUK PV 35057 (Figs. 6A-B; Supp. Fig. 3). The anterior and posterior ceratohyals are similar in size, and join one another along a subvertical junction. There is no obvious suturing between the two bones, and together they form a lozenge-shaped structure. There is no visible foramen in the anterior ceratohyal. The posterior ceratohyal is an obtuse trapezoid in lateral view and bears a fossa on its posterodorsal margin onto which the interhyal articulates. The interhyal is hooked, such that the larger ventral margin is in contact with the posterior ceratohyal—with the most dorsal point being at the midsection of the element—before curving laterally to form a small articular head in contact with the symplectic. The anterior ceratohyal tapers anteriorly, forming a well-developed articular head situated at the tip of a short process. This condyle articulates with a deep cavity in the posterior margin of the ventral hypohyal. This posterior excavation gives the ventral hypohyal a hook-shaped profile in lateral view. Dorsally, the ventral hypohyal articulates with a nodular, dorsal hypohyal. There is no suturing between these bones, and the displacement of the dorsal hypohyals suggests the articulation between the hypohyals was cartilaginous. The shape of the posterior margin of the dorsal hypohyal corresponds to the anterodorsal margin of the anterior ceratohyal. This, combined with the presence of a clear joint between the hypohyals and ceratohyals, suggests the potential for flexion within the ventral hyoid arch, as evident in *Lampris* (Fig. 6D).

A combination of incomplete ossification and displacement of bones makes interpretation of the branchial skeleton uncertain (Supp. Fig. 3), so our description is abbreviated. There are three basibranchials, increasing in length anteriorly to posteriorly. Five rod-like ceratobranchials are preserved. The first four ceratobranchials are relatively robust and grooved ventrally, while the fifth ceratobranchial is conspicuously more gracile, and consists of a slender rod without a groove. The hypobranchials are approximately half the length of their associated ceratobranchials. The epibranchials are poorly preserved and disarticulated from life position, making precise identification difficult.

### Vertebral column and dorsal fin

Six nearly complete vertebrae are visible in tomograms of NMS 1864.6.9 (Fig. 7A). The neural spines form the neural arches ventrally and are unpaired distally. The neural spines are straight and angled posteriorly. There are a series of epineurals contacting the neural spines on the first two vertebrae, and the base of the neural arches on vertebrae 3–6. The preserved ribs are slender and are visibly angled. Their articular heads directly contact the vertebrae at the level of their centra. The vertebral centra bear no parapophyses or haemal arches. A straight, rod-like bone lays at an oblique angle anterior to the most anterior neural spine. We interpret it as the ventral extremity of a single supraneural. No other supraneural is visible—though it is unclear if this is taphonomic or biological. No dorsal fin pterygiophores are visible, presumably because their ventral extremities were dorsal to the area that is preserved in the specimen.

**Figure 7.**
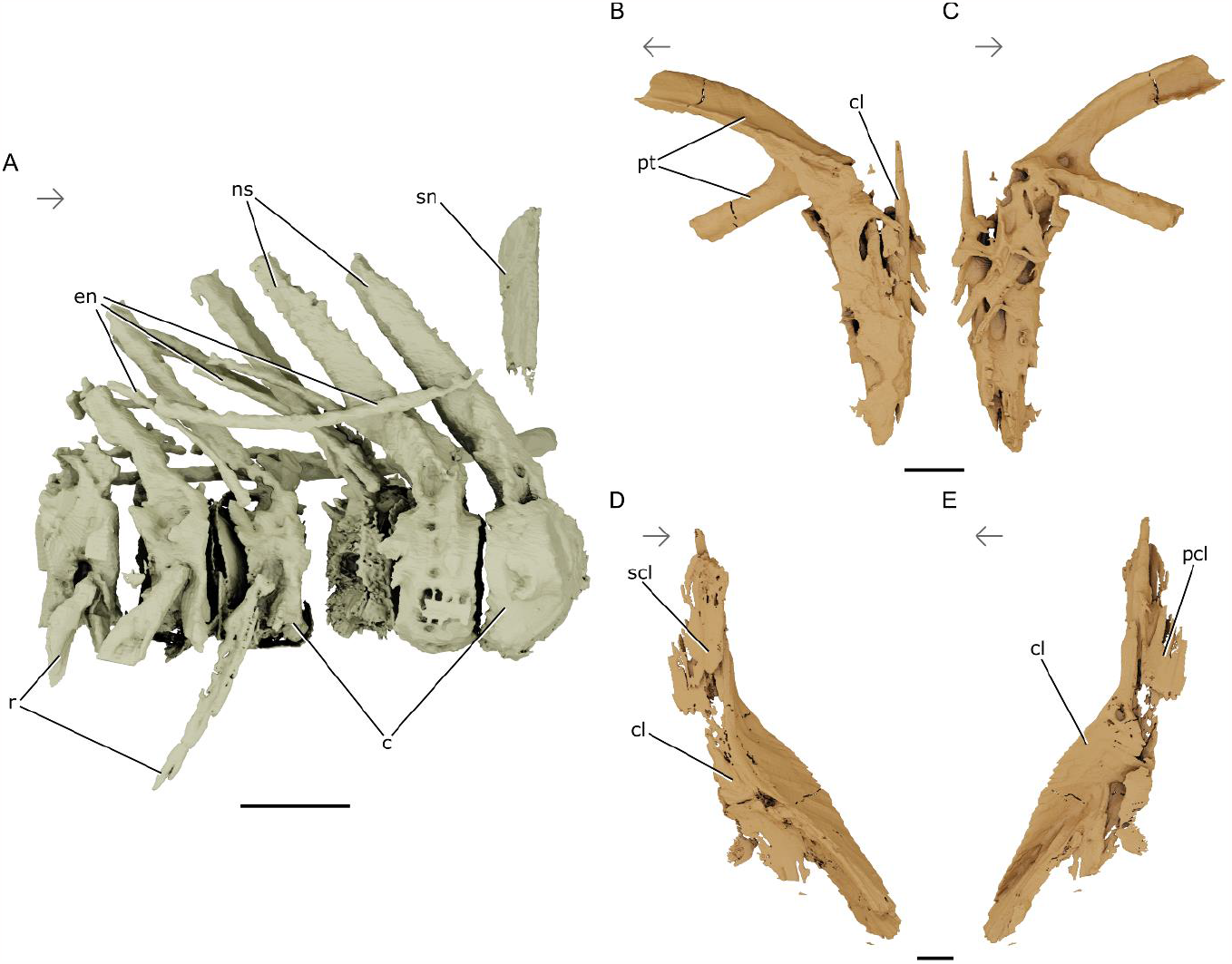
Postcranial elements of †*Whitephippus* cf. *tamensis*. NMS 1864.6.9, London Clay Formation, early Eocene (Ypresian), Southeast England, UK. Rendered μCT models showing the (**A**) vertebral elements in right lateral view; dorsal pectoral girdle elements in (**B**) lateral and (**C**) mesial views; ventral pectoral girdle elements in (**D**) lateral and (**E**) mesial views. Abbreviations: **c**, centra; **cl**, cleithrum; **en**, epineurals; **ns**, neural spines; **pcl**, postcleithrum; **pt**, posttemporal; **r**, ribs; **scl**, supracleithrum; **sn**, supraneural. Arrows indicate anatomical anterior. Scale bars represent 1 cm.

### Pectoral girdle and fin

NMS 1864.6.9 best preserves the dorsal portions of the pectoral girdle on the left side of the specimen, while the ventral portions of the pectoral girdle are best preserved on the right (Fig. 7B-E). The dorsal and ventral branches of the posttemporal extend the length of three vertebrae. The dorsal branch ends in a broad, flat head that articulates with the epioccipital. The ventral branch ends in a cylindrical head that articulates with the intercalar. Tomographic cross-sections of the posttemporal show highly mineralized struts along the long axis of the bone. The ventrolateral margin of the posttemporal articulates with a shattered supracleithrum whose shape cannot be adequately described. Mesial to the supracleithrum is the dorsal point of the cleithrum, which is sharply pointed and rises to the approximate height of the posttemporal bifurcation. Ventrally, the cleithrum comprises two flanges—one that continues laterally from the more dorsal portion, and another that penetrates mesially and bears an anteriorly-oriented convexity, such that the cleithrum appears as a chevron in coronal cross-section.

The pectoral fin is best preserved in NHMUK PV 35057, where the posterior half of the scapula, anterior portions of the coracoid, a complete series of four radials, and pectoral fin rays are present (Fig. 9). Additionally, fragments of the postcleithrum are visible between the chondral components of the girdle.

The scapula is broken along the scapular foramen on both the left and right side of the specimen, with a predominantly flat dorsal edge dropping nearly 90° to a swooping margin abutting the coracoid. At the anterodorsal margin of each break is an enlarged knob that forms an embayment for the propterygium (first pectoral radial). The scapular foramen appears to have been exceedingly large relative to the remainder of the pectoral skeleton: its diameter is more than half the height of the scapula. The coracoids are poorly preserved, with only the most dorsal portions articulated well enough to figure. The concave dorsal margin of the coracoid follows the convex ventral margin of the scapula, and its posterior border is similarly a continuation of the posterior border of the scapula.

The radials are displaced, making their identity difficult to ascertain. Despite this, four independent sizes of radials are evident, suggesting that there were four pairs in life. They appear to have extended posteroventrally past the margin of the scapulocoracoidal boundary, such that they would have sat along the cartilage between the scapula and coracoid in life. The anteriormost pectoral radial is stout— reaching a height less than half that of the fourth pectoral radial—and possesses a perforation / canal common to actinopterygian propterygia (Friedman, 2015; Jessen, 1972), which confirms its identity as the first pectoral radial. The remaining pectoral radials are approximately hourglass shaped and increase in size across the series— though taphonomic displacement from life position has moved a central radial posterior to the fourth (last) radial on the right side of the pectoral skeleton (Fig. 9).

Pectoral fin rays (lepidotrichia) begin at the most anterior boundary of the first pectoral radial and span the length of where the four radials would have sat in life. Each fin ray bears a club-like articular head proximally that attenuates distally before thickening at the junction of the paired lepidotrichia.

Portions of the postcleithrum are visible mesial to the pectoral fin rays, and whether it formed one or two separate ossifications in life is unclear. They are anteroposteriorly elongate at their dorsal extremities and the ventral portions of the postcleithral shaft approximate a triangle in coronal cross-section.

### Squamation

Squamation is preserved over the pectoral girdle of NHMUK PV P 6479 (Fig. 9), and posterior to the skull of NMS 1864.6.9 (Fig. 1). In both cases the scales are large, cycloid, and slightly overlapping.

## DISCUSSION

### †*Whitephippus* as a lampriform

In his original description of the †*Whitephippus* specimens studied here, Woodward (1901) attributed them to †*Laparon alticeps* and to Ephippidae. In Casier’s (1966) monograph on London Clay fossils, †*Whitephippus* is recognized as a new genus and species due to numerous differences with †*Laparon*. The latter, known by a single 3D-preserved specimen that has not been redescribed in details since, is still referred to as an ephippid in recent publications (Friedman et al., 2016). Casier (1966) does not call into question the attribution of †*Whitephippus* to Ephippidae, although he calls it a “less specialized ephippid” than †*Laparon* and *Ephippus*, and notes that the pectoral girdle is similar to that of *Lampris* (see below). Broad phenetic similarities between †*Whitephippus* and ephippids include a large, rectangular lachrymal, frontals contributing to the supraoccipital crest, and the anterior margins of the frontals curving dorsally at the midline. None of these characters are exclusive to ephippids. In contrast, phenetic dissimilarities between †*Whitephippus* and total group ephippids include the large, plate-like opercles in †*Whitephippus* (Figs. 1, 2) while it is smaller and more angular in ephippids (Cavalluzzi, 2000), the horizontal orientation of the pectoral fin in †*Whitephippus* (Figs. 1B; 9) while it is nearly vertical in ephippids, and the single-headed articular condyle of the hyomandibula while it is two-headed in ephippids (Cavalluzzi, 2000).

Anatomical studies strongly support lampriform monophyly (Davesne et al., 2014, 2016; Delbarre et al., 2016; Oelschläger, 1983; Olney et al., 1993; Wiley et al., 1998) with a series of morphological characters that, although not necessarily unique to the clade, are diagnostic when found in combination. Among these characters, all of those that are observable in †*Whitephippus* correspond to the lampriform state. These include synapomorphies of total group Lampriformes or of smaller clades encompassing parts of the lampriform stem (Davesne et al., 2014; Delbarre et al., 2016): (1) the antorbital is absent (Figs. 1–2), (2) the supraneural inserts anterior to the first neural spine (Fig. 7A), (3) the median ethmoid is partially posterior to the lateral ethmoids (Figs. 3A, D; 4A, D, H), (4) the frontals participate to the anterior portion of the sagittal crest (Figs. 3A, E; 4A, B, H); and synapomorphies of crown Lampriformes (Davesne et al., 2014, 2016): (1) the palatine lack an anterior process that articulates with the upper jaw (Fig. 5), (2) the frontals form an anterior cavity or ‘vault’ (Figs. 3A, C, E, I; 4B, D, F), and (3) the anterior extremity of the anterior ceratohyal forms a condyle that articulates with the ventral hypohyal (Fig. 6). Moreover, †*Whitephippus* shows character states that are plesiomorphic for acanthomorphs as a whole and differ in other clades such as percomorphs (including ephippids). These include the single-headed condyle of the hyomandibula (Fig. 5C, F, H), the presence of an orbitosphenoid (Figs. 3A, G, 4A, H), and the supraoccipital not contributing to the dorsal roof of the foramen magnum (Fig. 3F, L, 4E, I, Supp. Fig. 1) (Davesne et al., 2016; Johnson & Patterson, 1993; Olney et al., 1993).

The attribution of †*Whitephippus* to Lampriformes is then strongly supported by the available anatomical information, in contradiction with Woodward’s (1901) and Casier’s (1966) interpretation as an ephippid.

### Position within Lampriformes

Lampriform intrarelationships are largely congruent between phylogenetic studies based on morphological characters: *Lampris* forms a clade with the elongate Taeniosomi, to the exclusion of veliferids (Davesne et al., 2014, 2016; Delbarre et al., 2016; Olney et al., 1993; Wiley et al., 1998). Available evidence suggests that †*Whitephippus* is associated with this nested lampriform group.

The *Lampris* + Taeniosomi clade is notably characterised by a pectoral fin inserting horizontally and supported by three autogenous radials (Fig. 8E, F), the anterior-most being fused to the scapula (Olney et al., 1993). The pectoral fin of †*Whitephippus* inserts horizontally in the only specimen that preserves it (Fig. 1B, 9). However, the anterior-most pectoral fin radial is not fused to the scapula (Fig. 9), corresponding to the plesiomorphic lampriform arrangement (Fig. 8C, D). *Lampris* and taeniosomes also lack a dorsal foramen in the ceratohyal (Oelschläger, 1983), with †*Whitephippus* showing this same condition (Fig. 6A, B, D). In *Lampris* and taeniosomes, the infraorbital series consists in the lacrimal only (Oelschläger, 1983); this is also the case in †*Whitephippus*, but it is possible, although unlikely, that it is due to incomplete preservation (Figs. 1A, D, 2B). At the same time, typical taeniosome characters (Olney et al., 1993) are lacking in †*Whitephippus*: the neural spines of the anterior vertebrae are inclined posteriorly, rather than anteriorly like in taeniosomes (Figs. 7A, 8A, B), and there is at least one supraneural bone (supraneurals are absent in taeniosomes).

**Figure 8.**
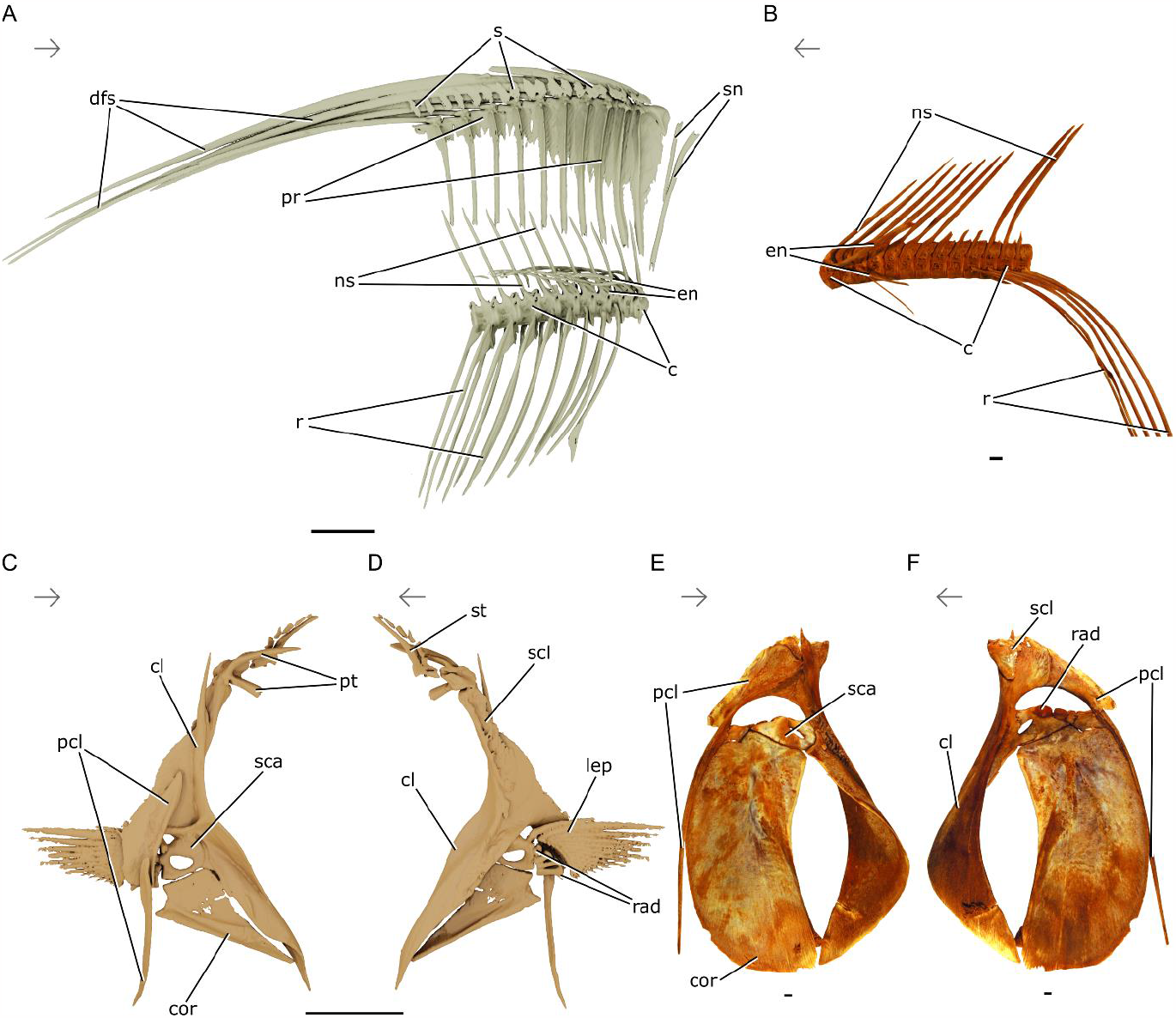
Postcranial elements of relevant lampriforms. *Velifer hypselopterus* UMMZ 220456, rendered μCT model of anterior vertebral elements in (**A**) right lateral view; *Lampris* cf. *guttatus* AMNH 79669SD, photograph of anterior vertebral elements in (**B**) lateral view; rendered μCT models of pectoral girdle of *V. hypselopterus* UMMZ 220456 in (**C**) lateral and (**D**) mesial views; photographs of pectoral girdle of *L. guttatus* AMNH 79669SD in (**E**) lateral and (**F**) mesial views. Abbreviations: **c**, centra; **cl**, cleithrum; **cor**, coracoid; **dfs**, dorsal fin spines; **en**, epineurals; **lep**, lepidotrichia; **ns**, neural spines; **pcl**, postcleithrum; **pr**, proximal radials; **r**, ribs; **rad**, radials; **s**, scales; **sca**, scapula; **scl**, supracleithrum; **sn**, supraneurals; **st**, supratemporal. Arrows indicate anatomical anterior. Scale bars represent 1 cm.

**Figure 9.**
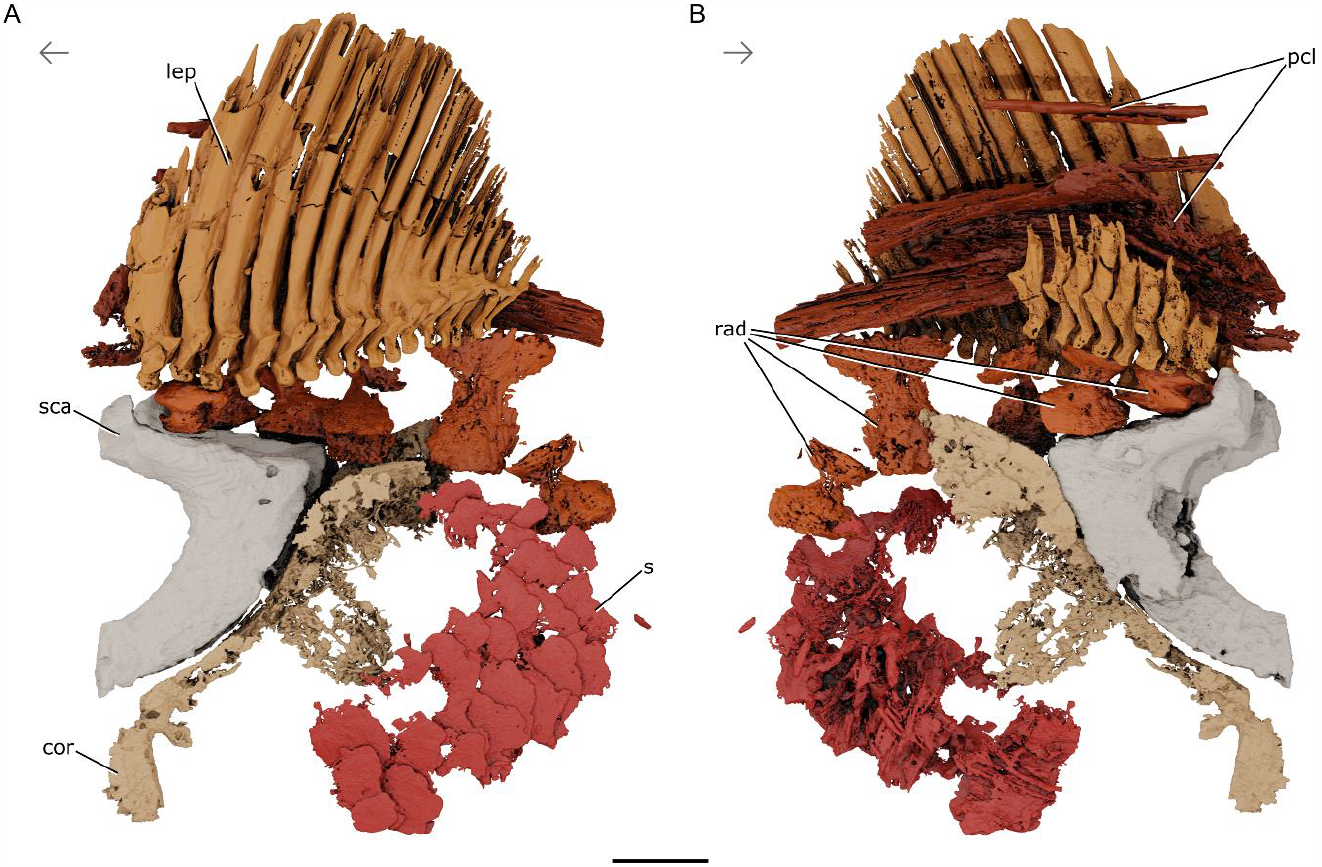
Pectoral fin skeleton of †*Whitephippus tamensis* holotype. NHMUK PV P 6479, London Clay Formation, early Eocene (Ypresian), Southeast England, UK. Rendered μCT models showing the (**A**) lateral and (**B**) medial views of the pectoral fin elements. Abbreviations: **cor**, coracoid; **lep**, lepidotrichia; **pcl**, postcleithrum; **rad**, radials; **s**, scales; **sca**, scapula. Arrows indicate anatomical anterior. Scale bar represents 1 cm.

Finally, two anatomical features reminiscent of extant *Lampris* are found in †*Whitephippus*: (1) the exoccipital condyles have dorsoventrally expanded, kidney-shaped posterior facets (Figs. 3F, 4I; Supp. Fig. 1) that define the lateral wall of the foramen magnum (Oelschläger, 1983; Olney et al., 1993) and (2) the four anterior-most vertebrae lack vertebral parapophyses (Figs. 7A; 8B) —there are developed parapophyses on all vertebrae in veliferids (Fig. 8A).

The available †*Whitephippus* material then shows a unique combination of character states, either shared with the *Lampris* + Taeniosomi clade (pectoral fin inserting horizontally, only one bone in the infraorbital series, no foramen in the anterior ceratohyal) or more specifically with *Lampris* (dorsally-expanded exoccipital condyles, lack of parapophyses on the anterior vertebrae). This would support that †*Whitephippus* is a member of the *Lampris* + Taeniosomi clade, and possibly an early member of Lampridae. One character however (the anterior-most pectoral fin radial not fused to the scapula), contradicts this attribution. The exact phylogenetic position of †*Whitephippus* within lampriformes is therefore best considered uncertain. A comprehensive phylogenetic analysis of the group including its well-preserved and anatomically disparate Paleogene fossil members—sometimes collectively termed “veliferoids”—is sorely needed.

It is to be noted that the few molecular studies in which veliferids are included tend to support a different set of deep divergences within lampriforms. Analyses of the 12S mtDNA (Wiley et al., 1998), a combination of six mitochondrial and nuclear markers (T. Grande et al., 2013), and complete mitogenomes (Wang et al., 2023) have all recovered a *Lampris* + veliferid clade sister to taeniosomes, contradicting morphological analyses. Future discussions of the phylogenetic position of †*Whitephippus* (and fossil lampriforms in general) will need to acknowledge this possible topology.

### Variation within †*Whitephippus* specimens

Our figured specimens identified as *†Whitephippus* show proportional and phenotypic differences, suggesting that there may be more than one species currently attributed to *†Whitephippus* in the London Clay formation. Variation includes parietal shape (more angular posteriorly in NMS 1864.6.9 (Fig. 3I) than in NHMUK PV P 35057 (Fig. 3C), a more rounded exoccipital condyle in NMS 1864.6.9 (Fig. 3L) than in NHMUK PV P 35057 (Fig. 3F), orbitosphenoid oriented moreso anteriorly in NMS 1864.6.9 (Fig. 3G), while in NHMUK PV P 35057 the orbitosphenoid is oriented moreso ventrally (Fig. 3A) (though this could be taphonomic), and a distinctly lacking basisphenoid pedicel in NHMUK PV P 35057 (Fig. 3A), whereas it is clearly present in NMS 1864.6.9 (Fig. 3G). Owing to the thin nature of the basisphenoid pedicel, it may have been broken or otherwise lost during preservation in NHMUK PV P 35057. Casier (1966) also signals differences between NHMUK PV P 35057 and the holotype NHMUK PV P 6479, particularly at the level of the ethmoid region, with the lateral ethmoids presenting a different ornamentation in lateral view (Fig. 1A-D) These clear differences warrant further examination of these specimens, though this is outside the scope of the present work.

### Lampriform diversity in the Palaeogene

The lampriform lineage extends to the Upper Cretaceous, with the †pharmacichthyids of the Cenomanian of Lebanon, the Cenomanian–Campanian †’aipichthyoids’ and †*Nardovelifer* from the Campanian of Italy (Davesne et al., 2014, 2016; Delbarre et al., 2016). However, the oldest members of the lampriform crown group are Cenozoic in age. *Incertae sedis* (‘veliferoid’, e.g. deep-bodied) lampriforms from Paleocene deposits in Scandinavia: the Danian København Limestone of Denmark and Southern Sweden yields †*Bathysoma* and cf. †*Palaeocentrotus* (Adolfssen et al., 2017; Davis, 1890), with the former genus also known from an erratic boulder of the Thanetian Lellinge Greensand (Bonde & Leal, 2017; Friedman et al., 2023). The fish-bearing horizon of the Danata Formation of Turkmenistan, which appears to correspond to the Paleocene–Eocene boundary, yields †*Danatinia* and †*Turkmene* (Bannikov, 1999; Daniltshenko, 1968). The Ypresian Fur Formation of Denmark yields the enigmatic †*Palaeocentrotus* plus an undescribed, †*Analectis*-like taxon (Bonde, 1966; Bonde et al., 2008). The earliest lampriforms that can be confidently classified within modern families are two veliferids (†*Veronavelifer* and †*Wettonius*) from the late Ypresian of Bolca, Italy (Bannikov, 1990, 2014; Carnevale et al., 2014; Carnevale & Bannikov, 2018)–although Near et al. (2013) have argued that †*Turkmene* is a lamprid, without providing a detailed justification. Bolca also yields two taxa historically described as lampriforms: the enigmatic ‘*Pegasus*’ (Carnevale et al., 2014; Carnevale & Bannikov, 2018) and †*Bajaichthys* (Bannikov, 2014; Sorbini & Bottura, 1988), the latter now regarded as a morphologically distinctive zeiform (Davesne et al., 2017). The Paleogene lampriform fossil record includes a variety of taeniosomes, all assigned to Lophotidae: †*Eolophotes* from the Lutetian of Georgia (Daniltshenko, 1980), †*Protolophotus* and †*Babelichthys* from the late Eocene of Iran (Arambourg, 1943; Davesne, 2017; Walters, 1957) and †*Oligolophotes* from the early Oligocene of Russia (Bannikov, 1999). These Russian deposits also yield the *incertae sedis* deep-bodied genera †*Analectis* and †*Natgeosocus* (Bannikov, 2014; Daniltshenko, 1980), with strata of comparable age in Germany yielding the veliferid †*Oechsleria* (Micklich & Bannikov, 2023). Unambiguous Lampridae are only known in the fossil record by the large-bodied †*Megalampris* from the Chattian (late Oligocene) of New Zealand (Gottfried et al., 2006) and by numerous specimens of *Lampris* from the Miocene of California (David, 1943; Jordan & Gilbert, 1919). If †*Whitephippus* is a lamprid, it would then be the first definitive Eocene representative of the family.

### Ecological implications

Modern veliferid lampriforms are predominantly demersal and neritic, living at depths not exceeding 250 m (Heemstra, 1986). Their ecological habits, but also their maximum size of approximately 40 cm, differ notably from the much larger size (up to 160 cm in *Lampris* and 800 cm in *Regalecus*; Hawn and Collette, 2012; Roberts, 2012) and epipelagic to bathypelagic lifestyle of other lampriforms. Based on this phylogenetic and ecological distribution, it appears likely that the proposed clade formed by Lampridae and taeniosomes to the exclusion of veliferids could be characterised as a ‘pelagic clade’. The relatively small sized, deep bodied stem lampriforms from the Upper Cretaceous are reminiscent of veliferids and presumably had similar ecological preferences as well (Delbarre et al., 2016). In contrast, most lampriforms from the late Paleocene and early Eocene are found in formations reflecting a more open-ocean environment, such as the Thanetian–Ypresian Danata Formation of Turkmenistan and the early Ypresian Fur Formation of Denmark (Bonde et al., 2008; Schrøder et al., 2022a, 2022b), to the exception of the veliferids from the late Ypresian Bolca Lagerstätte which is reconstructed as a shallow marine reef environment (Bellwood, 1996; Marramà et al., 2016). Similarly, the London Clay was deposited in an open-ocean, continental shelf paleoenvironment (King, 1981, 1984), which is reflected by its teleost fauna that is mostly (but not exclusively, e.g., Acipenseridae, Sparidae) composed of representatives of modern pelagic groups (Friedman et al., 2016). †*Whitephippus* is a putative member of the Lampris + Taeniosomi clade of pelagic lampriforms, and is recorded exclusively from the open marine paleoenvironment of the London Clay. It is then reasonable to infer that †*Whitephippus* was also an oceanic and probably pelagic animal, therefore documenting the transition in ecological preferences from proximal and demersal to oceanic and pelagic in lampriforms.

While the previous environment of the other pelagic acanthomorph clades is mostly unknown, their appearances in the fossil record are largely coordinated in time. The oldest scombrids are found in the Selandian of Angola (Dartevelle & Casier, 1949). The oldest carangids (Bannikov, 1985; Santini & Carnevale, 2015), billfishes (Fierstine, 2006; Monsch & Bannikov, 2011), and luvarids are all from the Paleocene–Eocene boundary of the Danata Formation. Deep-bodied lampriforms are also found at the latter horizon (Bannikov, 1999). The positions of these lampriform taxa are not clear, although some have suggested they might have affinities to the ‘pelagic clade’ (e.g., Near et al., 2013). Another open ocean fauna slightly older than the London Clay is found from the Fur Formation of the early Ypresian of Denmark. It preserves other *incertae sedis* lampriforms as well as fossils attributed to other modern pelagic families such as scombrids, carangids, polymixiids, stromateids and nomeids (Bonde, 1966; Bonde et al., 2008; Schrøder et al., 2022a, 2022b). Of all the aforementioned early Paleogene oceanic teleost faunas (to which a few other clay formations preserving fossils in three dimensions can be added; Friedman et al., 2016), the London Clay is by far the most diverse: it includes gadiforms (e.g., †*Rhinocephalus*), ophidiiforms (†*Ampheristus*), trichiurids (†*Eutrichiurides*), gempylids (†*Progempylus*), scombrids (e.g., †*Eocoelopoma*, †*Micrornatus*), carangids (e.g., †*Eothynnus*), xiphioids (e.g., †*Xiphiorhynchus*) and now lampriforms with †*Whitephippus*.

Molecular studies seem to confirm the origin and rapid diversification of pelagic acanthomorph clades in the early Paleogene (Harrington et al., 2016; Miya et al., 2013; Near et al., 2013), as suggested by the fossil record, even if older ages of divergence are sometimes estimated (Friedman et al., 2019; Santini et al., 2013; Santini & Carnevale, 2015; Santini & Sorenson, 2013). In particular, the taxa that occupy today the large-bodied pelagic predator ecological niche all seem to appear around the Paleocene–Eocene boundary. These taxa convergently share peculiar morphological and ecological traits that are also found in *Lampris*: relatively large body sizes, a crescentic caudal fin with the base of fin rays enveloping the caudal fin skeleton (Supp. Fig. 6), and a predatory behaviour that involves fast, active swimming over long distances and dives below the warm surface water. Billfishes, tunas and *Lampris* also independently developed a form of localised endothermy in the braincase, complemented in tunas and *Lampris* by a whole-body endothermy generated by axial and pectoral-fin red muscles, respectively (Dickson & Graham, 2004; Legendre & Davesne, 2020; Wegner et al., 2015). Whether †*Whitephippus* had the same kind of postcranial morphology is mostly unknown, but it might be possible to estimate if its metabolism was similar to that of modern *Lampris* by using bone histology as a proxy (Davesne et al., 2018).

The novel attribution of †*Whitephippus* to lampriforms may extend the taxon list of early pelagic fauna of the London Clay. If confidently attributed to the Lampridae total group, it reinforces the apparent observation that multiple lineages of acanthomorph teleosts simultaneously conquered the pelagic environment following the faunal depletion resulting from the Cretaceous–Paleogene mass extinction by separately acquiring similar morphological and physiological adaptations. Open ocean faunas from this critical time interval are few and far between (Argyriou & Davesne, 2021; El-Sayed et al., 2021; Friedman et al., 2016, 2023), and numerous taxa from the London Clay, but also from other early Paleogene open ocean faunas (e.g., from the Fur Formation), remain incompletely described (e.g. without knowledge of internal anatomy) and without a precise systematic attribution. An in-depth revision of these exceptional fossils is needed to achieve a better understanding of the evolutionary dynamics underlying acanthomorph teleost diversification and the establishment of modern marine faunas.

## ACKNOWLEDGEMENTS

We thank Stig Walsh (NMS), Emma Bernard and Zerina Johanson (NHMUK) for providing access to the fossil material, and Radford Arrindell (AMNH), Patrice Pruvost and Zora Gabsi (MNHN) and Randy Singer (UMMZ) for providing access to extant comparative material. Daniel Delbarre (University of Oxford) and Niels Bonde (University of Copenhagen) are thanked for their insightful discussions on the fossil specimens. We thank Carol Abraczinskas (University of Michigan Museum of Paleontology) for assistance in scanning line drawings.

Associate editor Jürgen Kriwet and two anonymous reviewers contributed to improve an earlier version of the manuscript. This study includes data produced in the CTEES facility at University of Michigan, supported by the Department of Earth and Environmental Sciences and the College of Literature, Science, and the Arts. This research was supported by the Natural Environment Research Council (grant NE/J022632/1 to MF), the University of Michigan Rackham Graduate School (Rackham Merit Fellowship to JVA), and the Alexander von Humboldt Foundation (Alexander von Humboldt-Stiftung to DD).

## DATA AVAILABILITY

The tomograms and surface files of segmented elements are available on MorphoSource (www.morphosource.org/projects/000561545). The segmented datasets are available on FigShare (URL XXXXX).

## AUTHOR CONTRIBUTIONS

DD and MF designed the project, DD, JVA and MF wrote the manuscript, DD, JVA, HTB and SG produced the data, JVA prepared the figures, all authors approved the final version of the manuscript. DD and JVA contributed equally to this work.

## SUPPLEMENTARY FILES

Supplementary_file_1.pdf

