## Supplementary file 1 for "Three-dimensional anatomy of the early Eocene *Whitephippus* (Teleostei: Lampriformes) documents parallel conquests of the pelagic environment by multiple teleost lineages"

### Three-dimensional anatomy of the early Eocene *Whitephippus* (Teleostei, Lampriformes) documents parallel conquests of the pelagic environment by multiple teleost lineages

DONALD DAVESNE,[1,2*] JAMES V. ANDREWS,[3,4,*] HERMIONE T. BECKETT,[2,5] SAM GILES,[6] and MATT FRIEDMAN[3,4]

[1]Museum für Naturkunde, Leibniz-Institut für Evolutions- und Biodiversitätsforschung, Invalidenstraße 43, 10115 Berlin, Germany,;

[2]Deparment of Earth Sciences, University of Oxford, South Parks Road, Oxford OX1 3AN, United Kingdom;

[3]Department of Earth & Environmental Sciences, University of Michigan, 1100 North University Avenue, Ann Arbor, Michigan 48109-1382, U.S.A.,;

[4]Museum of Paleontology, University of Michigan, 1105 North University Avenue, Ann Arbor, Michigan 48109-1382, U.S.A.;

[5]Department of Biology, King’s High School for Girls, Banbury Road, Warwick CV34 6YE, United Kingdom,;

[6]School of Geography Earth and Environmental Sciences, University of Birmingham, Birmingham B15 2TT, United Kingdom,

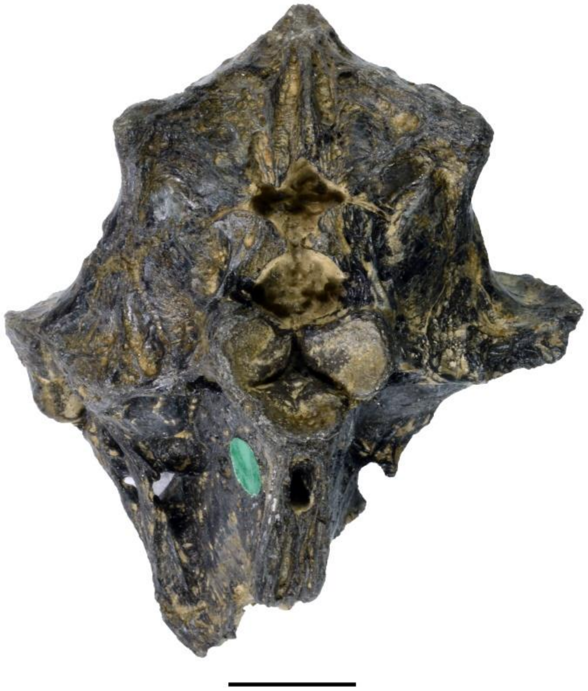

**Supplemental Figure 1.** Neurocranium of †*Whitephippus tamensis*, paratype NHMUK PV 41384 in posterior view. Scale bar represents 1 cm.

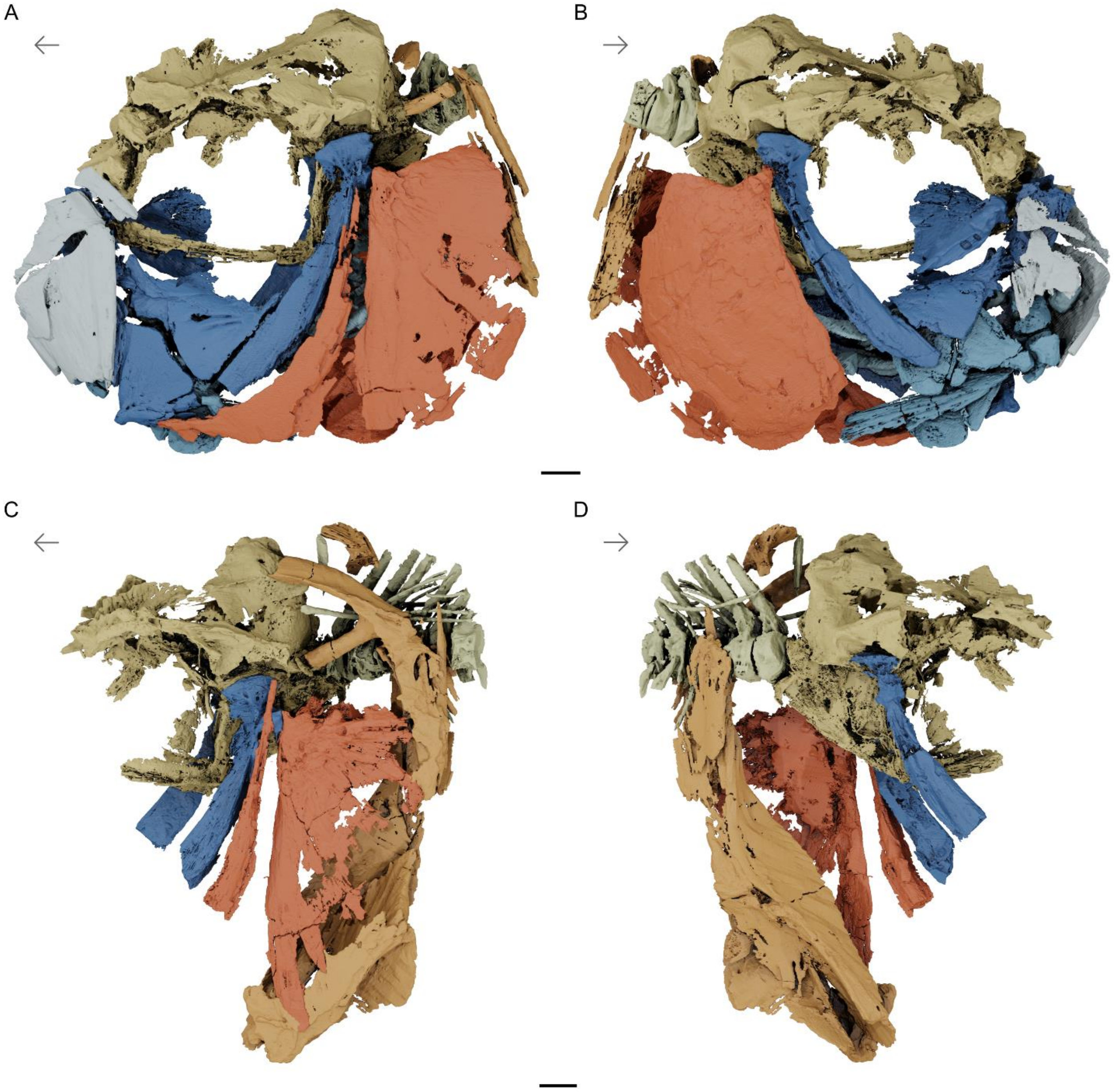


**Supplemental Figure 2.** Skull, vertebrae, and pectoral girdle of †*Whitephippus* cf. *tamensis*. London Clay Formation, early Eocene (Ypresian), Southeast England, UK. Rendered μCT models showing NHMUK PV P 35057 in (**A**) left and (**B**) right side views; NMS 1864.6.9 in (**C**) left and (**D**) right side views. Skeletal regions highlighted as follows: neurocranium (yellow), suspensorium (dark blue), jaws (platinum), opercles (sienna), ventral hyoid (blue), gill skeleton (light blue), pectoral girdle (orange), vertebral column (light yellow). Arrows indicate anatomical anterior. Scale bars represent 1 cm.

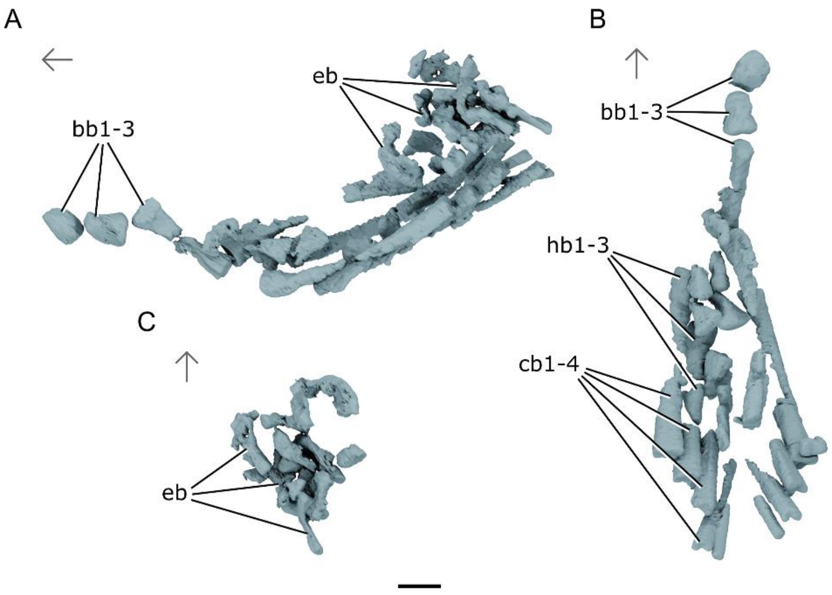


**Supplemental Figure 3.** Branchial skeleton of †*Whitephippus* cf. *tamensis* NHMUK PV P35057 in (**A**) lateral and (**B**) dorsal, and (**C**) ventral views. Abbreviations: **bb**, basibranchials; **cb**¸ ceratobranchials; **eb**, epibranchials; **hb**, hypobranchials. Arrows indicate anatomical anterior. Scale bar represents 1 cm.

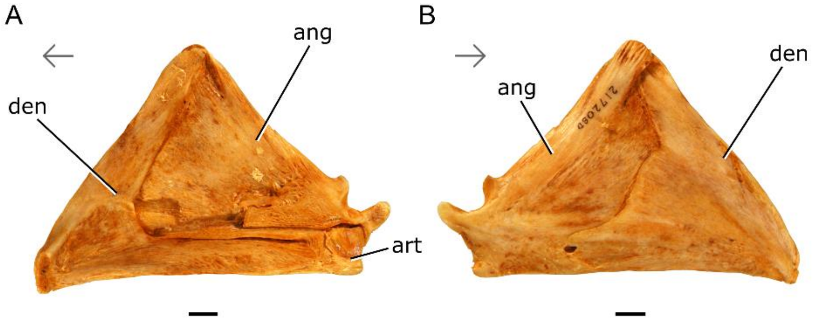


**Supplemental Figure 4.** Lower jaw of *Lampris* cf. *guttatus* AMNH 21720SD in (**A**) mesial and (**B**) lateral views. Arrows indicate anatomical anterior. Scale bars represent 1 cm.

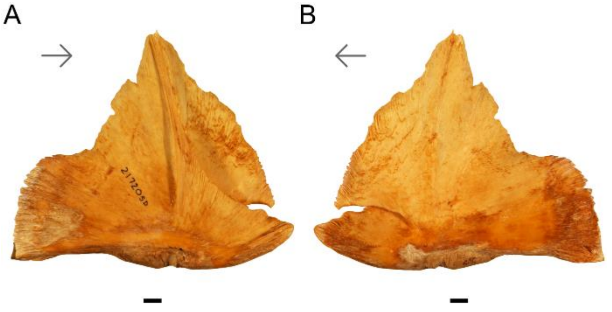


**Supplemental Figure 5.** Pelvic girdle of *Lampris* cf. *guttatus* AMNH 21720SD in (**A**) lateral and (**B**) mesial views. Arrows indicate anatomical anterior. Scale bars represent 1 cm.

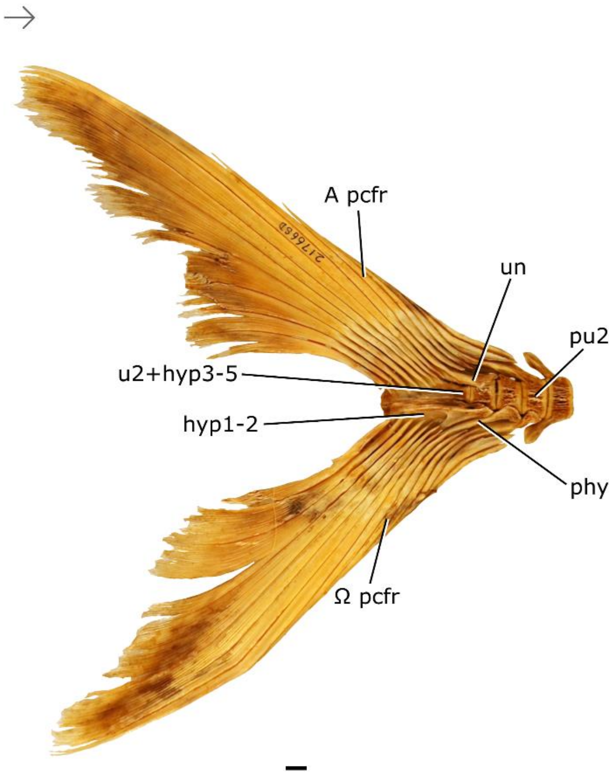


**Supplemental Figure 6.** Caudal skeleton of *Lampris* cf. *guttatus* AMNH 21766SD in lateral view. Abbreviations: **hyp**, hypural; **pcfr**, principal caudal fin ray (**A** = first; **Ω** = last); **phy**, parhypural; **pu**, preural; **u**, ural; **un**, uroneural. Arrow indicates anatomical anterior. Scale bar represents 1 cm.
